# Synchronised developmental partitioning of visual opsins across parental subgenomes in allopolyploid *Barbus barbus*

**DOI:** 10.64898/2026.10.12.758434

**Authors:** Zuzana Konvičková, Pau Balart-García, Demian Burguera, Veronika Truhlářová, Pavel Lepič, Zuzana Musilova

**Affiliations:** Department of Zoology, Faculty of Science, Charles University, Prague, Czech Republic; Faculty of Fisheries and Protection of Waters, South Bohemian Research Center of Aquaculture and Biodiversity of Hydrocenoses, University of South Bohemia in Ceske Budejovice, Czech Republic

## Abstract

Hybridisation and whole-genome duplication generate evolutionary novelty by combining and duplicating divergent parental genomes. Yet, how organisms differentially deploy duplicated gene repertoires across development remains poorly understood. The vertebrate visual system provides an excellent model for addressing this question because colour vision depends on the tightly regulated expression of opsin genes, which often undergo pronounced ontogenetic changes. Here, using comparative transcriptomics, we show that the allopolyploid common barbel (*Barbus barbus*) has evolved coordinated developmental partitioning of duplicated cone opsin genes. During ontogeny, the short-wavelength-sensitive opsins SWS1 and SWS2 switch expression between parental-subgenome ohnologs, while the middle-wavelength-sensitive RH2 switches between tandem paralogs, with all three transitions being synchronised. In contrast, the long-wavelength-sensitive LWS ohnologs from both parental genomes remain co-expressed throughout development. Single-nucleus RNA sequencing further revealed a strongly rod-dominated retina and largely conserved expression of phototransduction genes across recent ohnologs. The retention of all 14 visual opsin genes and their differential developmental deployment in common barbel highlights the value of independently evolved allopolyploid lineages for investigating the evolutionary consequences of genome duplication. More broadly, our findings show how hybridisation and genome duplication can enable divergent expression of parental gene copies, potentially contributing to sensory adaptation over evolutionary timescales.

## Main

Polyploidisation, or whole-genome duplication (WGD), duplicates the entire chromosome set and represents a major genomic perturbation that alters gene dosage and regulatory networks^1^. Following WGD, surviving lineages progressively return towards a diploid-like state, restoring genomic balance through gene loss, gene silencing, and divergence of retained gene copies^1,2^. WGD can arise through autopolyploidisation, involving genome duplication within a species, or allopolyploidisation which combines genome duplication with hybridisation between divergent progenitors^1^. In vertebrates, two ancient WGDs at the base of the lineage played a central role in morphological evolution, while a third, teleost-specific WGD likely contributed to the remarkable diversity of ray-finned fishes^3–5^. Despite growing interest in animal polyploidy, the evolutionary consequences of WGD in animals remain incompletely understood^6–8^.

Although lethal in mammals and birds, recent polyploidisation events are well documented in fishes, particularly among temperate freshwater taxa^6,9^. Recent WGDs provide a unique opportunity to investigate the early stages of genome evolution following polyploidisation, including processes that become obscured or undetectable after prolonged diploidisation of ancient polyploid genomes^10,11^. Cyprinidae sensu stricto (formerly Cyprininae; Cypriniformes) represents the largest vertebrate polyploid group, comprising more than 700 species on various ploidy levels that arose through at least 13 independent WGD events, predominantly allopolyploidisations^12^. Most studies of cyprinid polyploids have focused on two model species, common carp (*Cyprinus carpio*) and goldfish (*Carassius auratus*), which share the same WGD event^6,12^. The common barbel (*Barbus barbus*) belongs to the European tetraploid barbel clade, which originated through an independent allopolyploidisation event between 29.8 and 18.9 Mya^12^. This independent evolutionary origin makes the common barbel a valuable model for examining the fate and divergence of duplicated genes in a distinct cyprinid polyploid lineage.

The vertebrate retina is a light-sensitive neural tissue composed of photoreceptors (rods and cones) and interconnected neuronal and glial cell types^13^. Photoreceptors detect light through visual pigments consisting of opsins bound to retinal, with photon absorption initiating the phototransduction cascade and generating a signal that is subsequently processed by downstream retinal neurons and transmitted to the brain via the optic nerve^14,15^. Many components of the phototransduction cascade, including some visual opsins, arose in the ancient vertebrate WGDs^16^. Vertebrates possess five major visual opsin classes: rhodopsin (RH1) expressed in rods specialised for scotopic vision, and four cone opsins (SWS1, SWS2, RH2, and LWS) that can support colour discrimination under photopic conditions^17^. The spectral sensitivity of each visual pigment is determined by the opsin protein and its interaction with retinal^18^. Visual opsins therefore represent one of the best-resolved genotype–phenotype systems in vertebrates, in which defined molecular changes can produce predictable shifts in spectral sensitivity with direct consequences for visual function^18–20^.

Fishes develop externally in aquatic environments and often encounter distinct ecological conditions and visual demands throughout ontogeny. Consequently, their visual systems undergo substantial developmental remodelling, with retinal plasticity often persisting into adulthood^21–23^. Changes in opsin expression enable rapid tuning of photoreceptor spectral sensitivity without major alterations to retinal morphology. Developmental switches in opsin expression have been documented across diverse teleosts and often involve duplicated opsins within the same class^24–27^. This developmental partitioning is thought to promote the long-term retention of expanded opsin repertoires^28^. However, nearly all previous studies have examined opsins originating from individual gene duplication events rather than the evolutionary fate of entire visual gene repertoires following WGD. The common barbel provides a rare opportunity to investigate how duplicated opsin repertoires are retained, developmentally regulated, and functionally partitioned in a natural allopolyploid vertebrate whose larvae and adults occupy distinct ecological niches.

In this study, we investigate the visual system of the common barbel, an evolutionary tetraploid teleost. Using comparative transcriptomics across developmental stages and a single-nucleus RNA (snRNA-seq) atlas of the adult retina, we examine the consequences of polyploidisation for visual system evolution. We focus particularly on opsin genes retained from the recent WGD event, their evolutionary trajectories and expression patterns in the retina. We further localise opsin expression in the adult retina using fluorescence in situ hybridisation and assess the potential functional consequences of an expanded opsin repertoire for colour vision and spectral sensitivity.

## Results and Discussion

### Allopolyploidy retains an expanded and diversified visual opsin repertoire

The common barbel genome contains 14 visual opsin genes (seven from each subgenome), comprising ten cone opsins (SWS1 class: *sw1-A*, *sw1-B*; SWS2 class: *sw2-A*, *sw2-B*; RH2 class: *mw3-A*, *mw3-B*, *mw4-A*, *mw4-B*; LWS class: *lw1-A*, *lw1-B*) and four rod opsins (*rho-A*, *rho-B*, *rhol-A*, *rhol-B*) (Fig. 1A,B). This repertoire reflects an ancient allopolyploidisation event in the European tetraploid barbel lineage (29.8–18.9 Mya)^12^, superimposed on two earlier duplication events: (i) an ancestral duplication in clupeocephalans that gave rise to the rhodopsin-like (*rhol*) and rhodopsin (*rho*) lineages^29^, and (ii) a tandem duplication of RH2 resulting in the *mw3* and *mw4* paralogs. The *rho* and *rhol* copies are conserved across cyprinids and are therefore also present in the common carp and zebrafish. In contrast, the RH2 tandem duplication is absent from zebrafish but predates the independent allopolyploidisation events that gave rise to common barbel and common carp (Fig. 1A). Overall, the common barbel possesses approximately twice the average number of visual opsin genes found in teleosts^30,31^.

**Figure 1:**
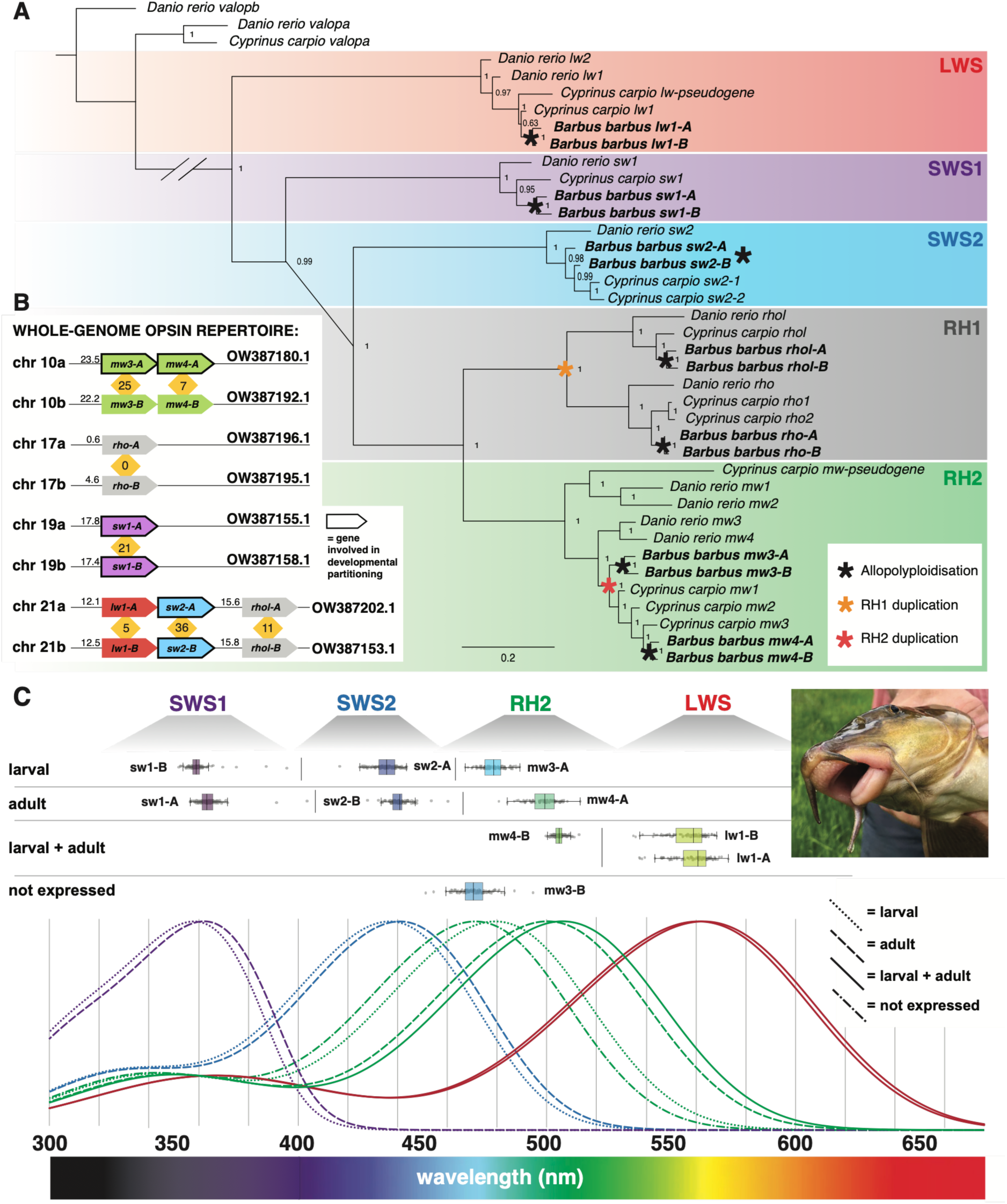
Diversity of the opsin gene repertoire in common barbel. (A) Phylogeny of the visual opsin genes in the common barbel (*Barbus barbus*), common carp (*Cyprinus carpio*), and zebrafish (*Danio rerio*). Bayesian majority-rule consensus tree, node labels indicating posterior probabilities. Vertebrate ancestral opsins from zebrafish and common carp were used as outgroups. Colours indicate opsin classes. Black asterisks mark the WGD event in the common barbel. The yellow asterisk indicates RH1 duplication event in the clupeocephalan ancestor that gave rise to a second rhodopsin gene (rhodopsin-like, *rhol*) in cyprinids, and the red asterisk denotes an RH2 (green-sensitive opsin) gene duplication in the ancestor of common barbel and common carp. (B) Positions of opsin genes on common barbel chromosomes based on the fBarBab1.1 genome assembly (NCBI accession GCF_936440315.1). Conversion between the chromosome nomenclature used in this study and the chromosome names and accession numbers in the assembly is provided in Supplementary Table S6. The “A” and “B” labels (e.g., 10a, 10b) distinguish chromosomes within a pair but do not consistently correspond to the two subgenomes. Approximate chromosomal positions are shown in megabases. Colours denote opsin classes as in A. Genes involved in developmental expression partitioning are highlighted with a black outline. Numbers in yellow indicate the number of amino acid substitutions between members of each ohnologous pair. (C) Predicted spectral sensitivities of cone opsins in the common barbel modelled using the OPTICS tool^33,34^, the whole-dataset model and amino-acid property encoding method (top), and absorbance curves of individual opsins based on their estimated absorption maxima (λ_max_) (bottom). Curve colours indicate opsin classes, while line styles denote developmental expression patterns as indicated in the legend. Curves were reconstructed using a template following Govardovskii et al. (2000)^35^.

All identified opsin coding sequences retained intact open reading frames, with no premature stop codons or frameshift mutations, allowing prediction of full-length proteins. Expression was detected for all opsin genes except *mw3-B* in the bulk RNA-seq data (see below), indicating that nearly all duplicated copies remain transcriptionally active and likely functional. The absence of detectable *mw3-B* expression despite an intact coding sequence suggests transcriptional silencing rather than pseudogenisation. Such patterns are common in teleosts, where expansion of individual opsin classes is frequently accompanied by transcriptional silencing or persistently low expression of specific gene duplicates (e.g., *RH2B* in *Pseudochromis fuscus*^32^, *LWS-2* and *LWS-3* in *Lucania goodei*^24^).

Opsin ohnologs differed by 0 (rho) to 36 (sw2) amino acid substitutions (Fig. 1B), indicating variable levels of sequence divergence among duplicated gene pairs. To assess potential functional consequences of this divergence, we predicted peak spectral sensitivity (λ_max_) using OPTICS^33,34^. The predictions indicated modest but consistent spectral differentiation among most cone opsin ohnologs, with estimated λ_max_ shifts of 4.2 nm for SWS1, 5.2 nm for SWS2, 7.9 nm for RH2 mw3, and 6.8 nm for RH2 mw4, whereas LWS ohnologs showed only minimal divergence (1.7 nm; Fig. 1C; Supplementary Table S2). In contrast, the two RH2 paralogs generated by the older tandem duplication (mw3 and mw4) exhibited pronounced spectral differentiation, with mw3 predicted to be blue-shifted by more than 20 nm relative to mw4.

### Duplicated opsins show synchronised developmental partitioning

We quantified visual opsin expression across eight developmental stages of the common barbel (5, 6, 8, 14, 21, 28 and 42 days post-fertilisation [dpf], and adults; Fig. 2, Supplementary Fig. S1D, Supplementary Tables S1 and S3). We identified synchronised ontogenetic switches between larval- and adult-expressed genes in three cone opsin classes (SWS1, SWS2, and RH2), with larval- and adult-expressed genes reaching similar expression levels at approximately 14 dpf (Fig. 2C). Notably, this developmental transition involved different categories of gene duplicates in different opsin classes. In the two short-wavelength-sensitive classes (SWS1 and SWS2), expression switched between ohnologs retained from the recent allopolyploidisation event, with *sw1-B* and *sw2-A* predominating during the larval stage and *sw1-A* and *sw2-B* in adults. For RH2, expression was shaped by two layers of gene duplication: developmental switching occurred between the earlier tandem duplicates *mw3-A* and *mw4-A*, whereas the corresponding more recent ohnologs showed asymmetric expression patterns, with *mw4-B* constitutively expressed and *mw3-B* remaining silent throughout ontogeny. In contrast, expression of LWS and rod opsins (*rho* and *rhol*) remained stable once established, with no evidence of expression partitioning between ohnologs or older tandem duplicates. Developmental partitioning is therefore selectively deployed within the visual opsin repertoire rather than a general feature of duplicated opsin genes. Although developmental switches between opsin paralogs belonging to different opsin classes^36^ or within the same opsin class^24–27^ are widespread among teleosts^19^, the synchronised partitioning of two SWS ohnolog pairs, together with the additional RH2 switch, distinguishes the common barbel from previously described systems.

**Figure 2:**
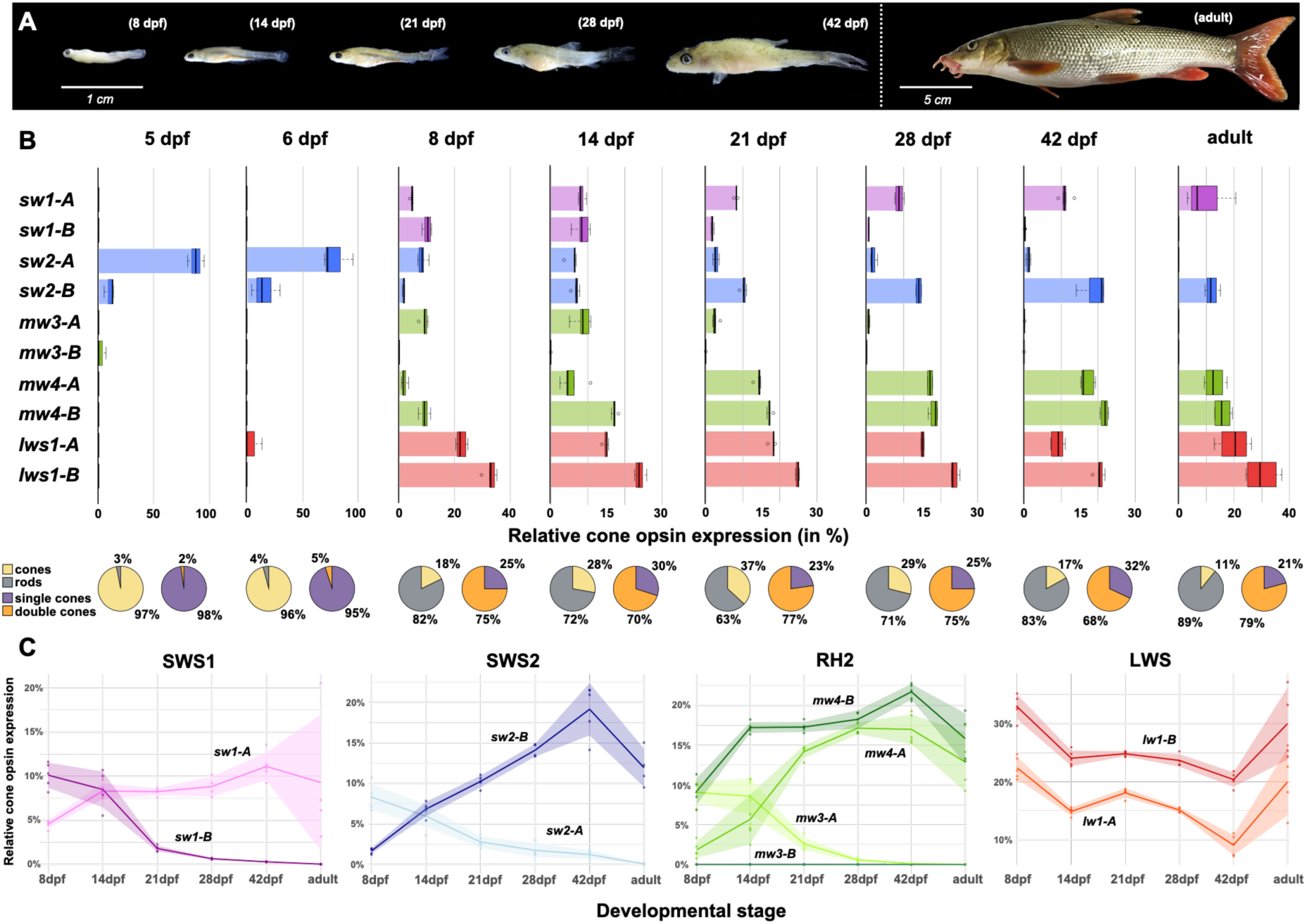
Development of the visual system in common barbel. (A) Photographs of developmental series of the common barbel. (B) Developmental series of opsin expression in the common barbel based on bulk retinal transcriptomes. Relative cone opsin expression is shown in box plots (top), with each opsin represented as a proportion of total cone opsin expression. Colours indicate cone opsin classes. The “-A” and “-B” suffixes denote the respective ohnologs but do not consistently correspond to the two subgenomes. Note the different x-axis scales. Pie charts (bottom) show the relative proportions of cone versus rod opsin expression and single-cone (SWS1 and SWS2) versus double-cone (RH2 and LWS) opsin expression, respectively. (C) Developmental changes in the opsin gene expression within individual cone opsin classes. Lines represent mean expression levels, while ribbons show standard deviations. A coordinated developmental expression switch occurs in the SWS1, SWS2, and RH2 opsin classes.

The developmental expression switches were accompanied by corresponding differences in predicted spectral sensitivity. The opsin classes exhibiting ontogenetic shifts (SWS1, SWS2, and RH2) also showed the greatest divergence in predicted λ_max_, whereas the constitutively expressed LWS and RH1 opsins remained spectrally similar. In both SWS classes, the larval-expressed ohnolog was predicted to be shifted towards shorter wavelengths relative to its adult counterpart (Fig. 1C). An even greater spectral difference was observed within RH2, where the larval-expressed mw3-A was predicted to absorb substantially shorter wavelengths (478.1 ± 4.8 nm) than both the adult-expressed mw4-A (498.4 ± 5.6 nm) and the constitutively expressed mw4-B (505.2 ± 5.2 nm). Consequently, the larval retina is predicted to be spectrally tuned towards shorter wavelengths than the adult retina. Moreover, the >20 nm separation between the strongly blue-shifted larval mw3-A and constitutively expressed mw4-B could broaden the spectral sensitivity of RH2 cones if these opsins were co-expressed within the same photoreceptors during larval development.

Shorter-wavelength spectral tuning in younger life stages has been documented across ray-finned fishes^37^. This developmental shift is generally interpreted as an adaptation to changes in habitat use and prey preference during ontogeny, with enhanced short-wavelength sensitivity potentially facilitating the detection of planktonic prey during early life^37–39^. Although the adaptive significance of the short-wavelength-shifted visual system in common barbel larvae remains to be tested, our findings are consistent with this general pattern. Expression of all cone opsin classes was established by 8 dpf, coinciding with the onset of exogenous feeding on plankton in the water column^40,41^. When juveniles reach a total length of approximately 20 mm, they transition to a benthic lifestyle and progressively acquire the morphology and feeding behaviour characteristic of crepuscular adult bottom feeders^40,42^. Notably, the developmental switch in opsin expression precedes the ecological transition, suggesting that visual system remodelling may precede, and potentially facilitate, subsequent shifts in habitat use and feeding ecology rather than merely responding to them. Opsin expression shifts preceding habitat transitions have been reported in other teleosts, such as the fangtooth (*Anoplogaster cornuta*)^43^, dusky dottyback (*Pseudochromis fuscus*)^32^, Atlantic tarpon (*Megalops atlanticus*)^44^ and European eel (*Anguilla anguilla*)^45^, suggesting that this may be a more general feature of visual development.

To our knowledge, the common barbel represents the first documented example of synchronised developmental partitioning of duplicated opsins involving both WGD-derived ohnologs and older paralogs across multiple cone opsin classes. Comparable ontogenetic analyses in other vertebrate lineages that have experienced additional WGDs, including other cyprinid allopolyploids, will be valuable for determining how general this pattern is. Our findings indicate that allopolyploidisation can be followed by synchronised developmental regulatory divergence among duplicated visual genes, providing a potential mechanism for the long-term retention and functional diversification of expanded opsin repertoires.

### Single-nucleus profiling reveals conserved retinal cell types in the adult common barbel

To investigate the expression of gene duplicates at the level of individual retinal cell types, we generated snRNA-seq datasets from two adult common barbel retinae (Supplementary Table S1). The datasets differed in whether nuclei were sorted prior to library preparation, allowing us to evaluate whether sorting affected dataset composition. After quality control, Sample 1 contained 4,670 nuclei and Sample 2 contained 5,888 nuclei with comparable levels of gene detection. Rod photoreceptors predominated in both datasets, comprising 89.2% and 82.2% of nuclei in Samples 1 and 2, respectively (Fig. 3A,B). Although the unsorted sample contained a slightly higher proportion of non-rod cell types, all annotated retinal cell types were recovered in both datasets (Supplementary Fig. S1B), indicating that nuclei sorting did not substantially alter overall cell-type representation.

**Figure 3:**
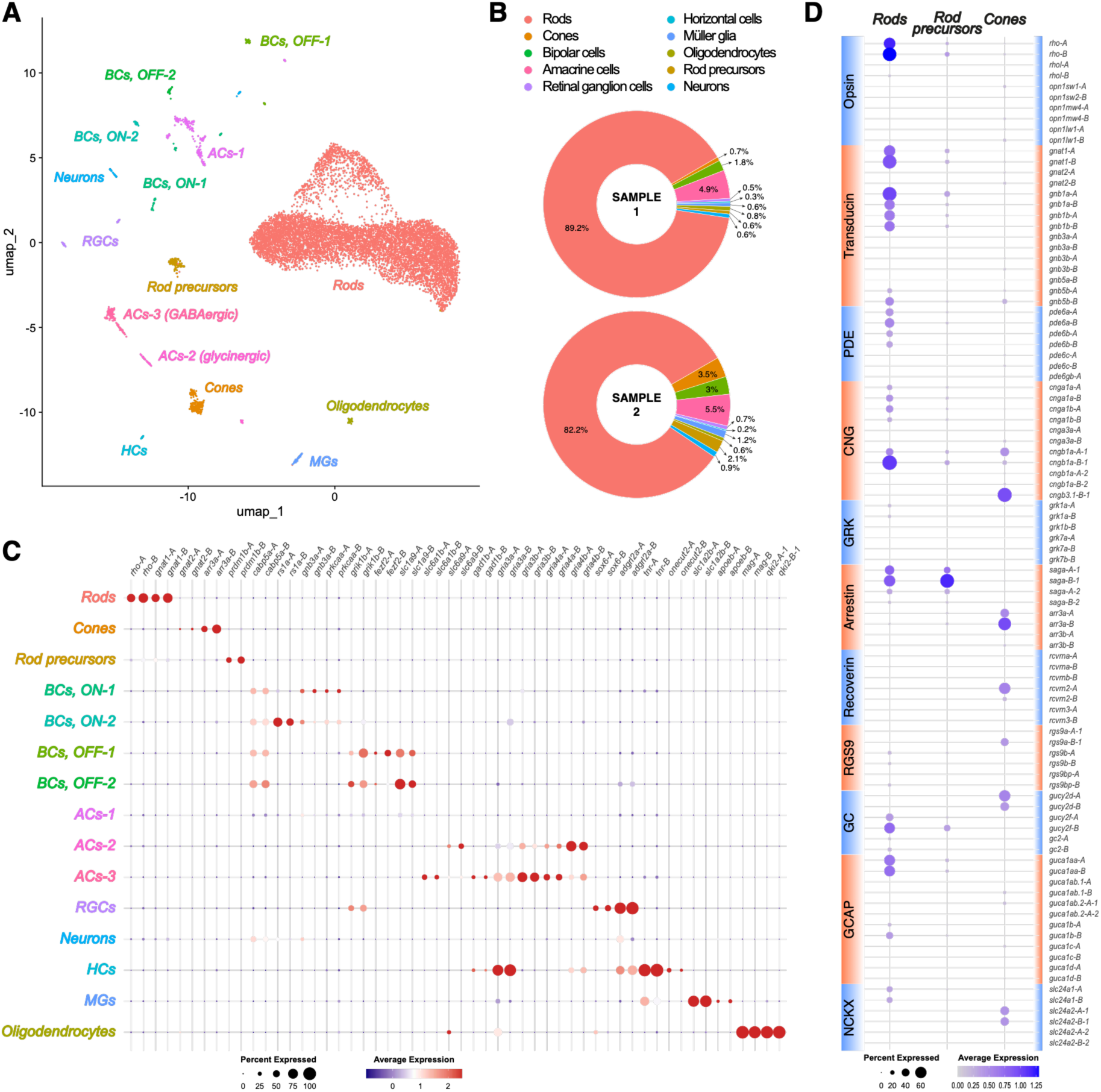
Single-nucleus characterisation of the common barbel retina. (A) UMAP visualisation of the integrated snRNA-seq dataset, with clusters annotated by retinal cell type. (B) Donut charts showing the proportions of major retinal cell types recovered for two common barbel samples profiled by snRNA-seq: Sample 1 (nuclei sorted prior library construction) and Sample 2 (unsorted nuclei). (C) Dot plot showing expression of representative marker genes enriched in individual cell type populations. Dot size indicates the percentage of cells expressing each gene, and dot colour gradient (red to indigo) represents scaled average expression levels. (D) Dot plot showing expression of phototransduction cascade genes across rod, rod precursor, and cone clusters. Ohnologs descending from recent allopolyploidisation distinguished by “-A” and “-B” suffixes. Dot size indicates the percentage of cells expressing each gene, and dot colour intensity represents normalised average expression levels.

Even though methodological differences preclude a direct quantitative comparison of cell-type proportions across studies, the proportion of rods observed in the common barbel was substantially higher than in published single-cell retinal datasets from other cyprinids^46–49^. For example, rods represented only 25.7% of retinal cells in goldfish^47^. This pronounced rod enrichment may reflect adaptation to the crepuscular, benthic lifestyle of adults, although the extent to which it reflects the cellular composition of the intact retina remains uncertain.

In total, we identified 18 transcriptionally distinct clusters representing 15 retinal cell types in the integrated filtered dataset (Supplementary Fig. S1A; Fig. 3A,C). These comprised all major retinal neuronal classes, together with Müller glia and oligodendrocytes. Rod photoreceptors resolved into four clusters, likely owing to their high abundance in the dataset, but their highly similar transcriptional profiles supported assignment to a single cell type (Supplementary Table S4). While several clusters could be assigned unambiguously based on highly specific marker genes (e.g., *rho* in rods and *mag* in oligodendrocytes), others exhibited overlapping expression profiles that precluded assignment based on a single diagnostic marker (see Methods).

Many marker genes defining major retinal cell types in the common barbel corresponded to those previously reported in goldfish, including *slc6a9* in glycinergic ACs-2, *gad1b* in GABAergic ACs-3, *tnr* in horizontal cells, and *apoeb* in Müller glia (Fig. 3C)^47^. An exception was observed for bipolar cells: whereas goldfish bipolar cells predominantly express *cabp5b*, common barbel bipolar cells express *cabp5a*, consistent with zebrafish^49^. Marker genes distinguishing ON and OFF bipolar cell subtypes likewise largely matched those reported in zebrafish^49^. Beyond the expected expression of *slc6a1* and *gad1b*, the GABAergic ACs-3 showed strong expression of multiple *gria* genes, suggesting substantial glutamatergic input. To our knowledge, this molecular signature has not been reported in previous single-cell studies of the teleost retina.

We identified several examples of expression partitioning between older paralogous marker genes. Consistent with previous observations in zebrafish, the *gnb3* paralogs displayed cell type-specific expression, with *gnb3a* enriched in ON bipolar cells and *gnb3b* in cone photoreceptors (Fig. 3C,D)^49^. Similarly, the *gria3* paralogs showed complementary expression patterns between horizontal cells (*gria3a*) and GABAergic ACs-3 (*gria3b*), whereas the *gria4* paralogs were partitioned among amacrine cell subtypes, with *gria4a* enriched in GABAergic ACs-3 and *gria4b* in glycinergic ACs-2. In contrast, the recent ohnologous marker gene pairs examined generally retained similar expression profiles across retinal cell types, although some showed clear cell type-specific partitioning (Fig. 3C). For example, *slc6a9-A* was predominantly expressed in oligodendrocytes, whereas *slc6a9-B* was enriched in glycinergic ACs-2. Similarly, *fezf2-B* was selectively expressed in one OFF bipolar cell subtype (BCs OFF-1) (Fig. 3C).

Together, these analyses establish a cellular atlas of the adult common barbel retina that recovers the major retinal cell types, broadly recapitulates established marker-gene expression patterns and provides a cellular framework for investigating the expression and partitioning of duplicated visual opsin, phototransduction and photoreceptor recovery genes in the adult retina.

### Expanded cone opsin repertoire is accommodated within four canonical cone types

The cone photoreceptor cluster was subsetted and reanalysed to investigate gene expression patterns among individual cone subtypes. Three cone clusters were identified based on differential gene expression and assigned to SWS1 (UV; 50 nuclei), RH2 (green; 40), and LWS (red; 138) cones (PR4, PR2, and PR1, respectively, following Baden et al. (2025)^50^; Fig. 4A). The top 10 differentially expressed genes were highly specific to their respective cone subtypes and included previously reported subtype-specific markers^49–51^, such as *thrb* (LWS), *arr3b* (SWS1/SWS2), *fat1a* (SWS1), *tgfa* (SWS1), *fibcd* (RH2), and *pcdh17* (RH2) (Fig. 4C; Supplementary Table S5). The expression of the corresponding opsin genes further supported these assignments (Fig. 4B). Although nuclei expressing SWS2 opsin were also detected, they did not form a distinct cluster (Fig. 4B), likely reflecting the limited number of cone nuclei in the dataset (228 in total) and sparse detection of opsin transcripts at the single-nucleus level.

**Figure 4:**
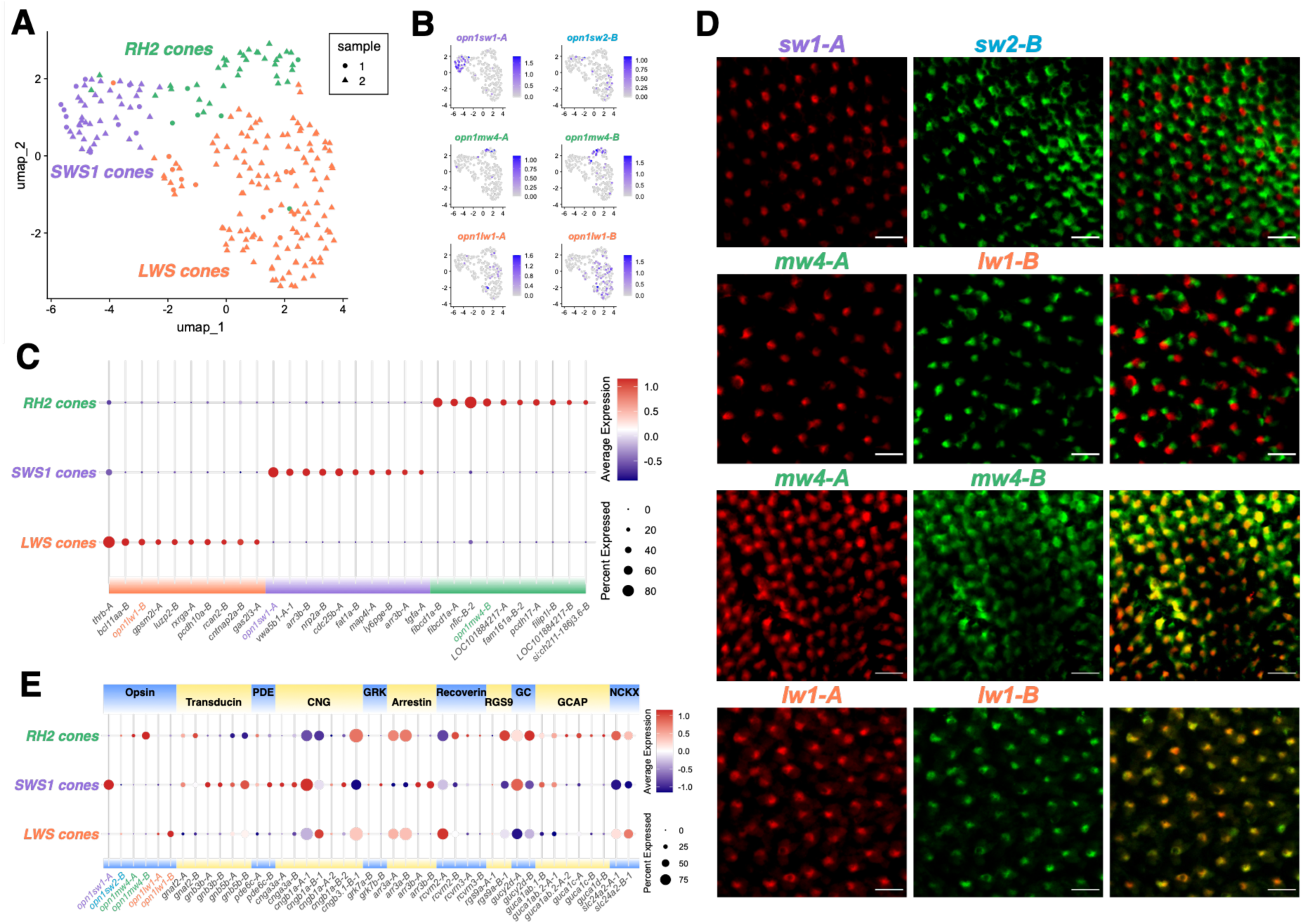
Cone photoreceptors in the common barbel. (A) UMAP embedding of cone photoreceptor nuclei from the integrated dataset, resolving three cone subtypes: SWS1, RH2, and LWS cones. Point shapes indicate sample of origin. (B) Feature plots showing normalised expression of all cone opsin genes expressed in the adult retina of the common barbel. Ohnologs are distinguished by “-A” and “-B” suffixes. Note that nuclei expressing the blue-sensitive opsin gene *opn1sw2-B* did not form a distinct cluster. (C) Dot plot showing expression of the top 10 marker genes for each cone cluster (see complete list in Supplementary Table S5). Dot size indicates the percentage of cells expressing each gene, and dot colour gradient (red to indigo) represents scaled average expression levels. (D) Fluorescence in situ hybridisation labelling of photoreceptors in the adult common barbel retina. Photoreceptors were labelled according to their opsin expression using opsin-specific probes. The first and second columns show the individual probe signals, while the third column presents the corresponding merged images. Opsins targeted by each probe are indicated above the images. The first row shows distinct expressions of *sw1-A* and *sw2-B* in separate single cones, which are arranged in a regular mosaic pattern. The second row demonstrates differential expression of *mw4-A* and *lw1-B* in opposing members of double-cone photoreceptors, which also exhibit regularity in their arrangement. The third and fourth rows illustrate co-expression of ohnologous opsins (*mw4-A* and mw4-B; *lw1-A* and *lw1-B*, respectively) within the same members of double cones. The scale bar represents 20 μm. (E) Dot plot showing expression of phototransduction cascade genes across cone types. Dot size indicates the percentage of cells expressing each gene, and dot colour gradient (red to indigo) represents scaled average expression levels.

We focused on the *mw4-A*/*mw4-B* and *lw1-A*/*lw1-B* ohnolog pairs, both of which are expressed in the adult retina and therefore represent candidates for expression specialisation. However, because RH2 and LWS cone clusters did not subdivide by ohnolog expression and opsin transcripts were sparsely detected, snRNA-seq alone could not determine whether these ohnologs are co-expressed within individual cones or partitioned among distinct photoreceptor populations. We therefore used fluorescence in situ hybridisation (FISH) to resolve the cellular expression patterns of these ohnologs and to characterise the spatial organisation of adult cones (Fig. 4D). This analysis revealed that SWS1 and SWS2 opsins (*sw1-A* and *sw2-B*, respectively) are expressed in distinct single cones, whereas RH2 and LWS opsins (*mw4-A* and *lw1-B*, respectively) are expressed in opposite members of double cones. Ohnolog-specific FISH further demonstrated co-expression of the *mw4-A*/*mw4-B* and *lw1-A*/*lw1-B* pairs within the corresponding double-cone members. Cone photoreceptors formed a square mosaic, with highly regular single cones and less ordered double cones (Fig. 4D). This finding contrasts with a previous study that reported an irregular cone arrangement in the common barbel^52^, but resembles the square mosaic described in many cyprinids, including goldfish^53–55^. The regular cone arrangement in the common barbel is consistent with the proposed role of cone mosaics in enhancing spatial resolution through uniform distribution of photoreceptor subtypes^56^.

Despite the expanded cone opsin repertoire and simultaneous expression of six cone opsin genes, only four canonical cone subtypes were identified in the adult retina. Unlike the zebrafish retina, where additional red-shifted cone populations have been identified from single-cell and histological data^49,57^, neither snRNA-seq nor FISH revealed evidence for additional cone populations with distinct expression profiles in the common barbel. Instead, RH2 and LWS opsin ohnologs were co-expressed within the corresponding cone types, at least in the dorsal retinal region examined by FISH. Although co-expression of opsins with divergent λ_max_ values can broaden photoreceptor spectral sensitivity or enable spectral tuning^19,58^, the functional consequences of RH2 and LWS ohnolog co-expression in adult double cones may be limited because their absorption maxima are very similar (Fig. 1C, Supplementary Table S2). Such effects could be more pronounced in larval RH2 cones, where the absorption maxima of the two simultaneously expressed opsins, mw3-A and mw4-A, differ by more than 20 nm, if these pigments are co-expressed.

Ohnologs retained following allopolyploidisation therefore appear to contribute to distinct expression strategies across cone types: in double cones, RH2 and LWS ohnologs are co-expressed, whereas expression of SWS1 and SWS2 ohnologs in single cones is developmentally partitioned, ultimately resulting in silencing of the larval-expressed ohnolog in the adult retina. Overall, our data suggest that the expanded opsin repertoire of the common barbel is partitioned primarily across ontogeny rather than among distinct photoreceptor populations. Thus, developmental regulation, rather than the emergence of novel photoreceptor populations, represents the major mechanism by which duplicated opsins have acquired distinct roles in the common barbel visual system.

### Phototransduction genes show limited expression divergence among recent ohnologs, whereas ancient paralogs exhibit expression partitioning

We complemented the opsin analysis by examining a broader set of phototransduction and photoreceptor recovery genes. Based on published literature^16,49^, we compiled a list of 103 genes, including opsins, of which 94 were detected in the snRNA-seq dataset. We analysed their expression across rods, rod precursors, and cones to assess whether ancestral paralogs and recent WGD-derived ohnologs showed evidence of expression divergence (Fig. 3D). Genes expressed in cone photoreceptors were further examined across the three cone subtypes (Fig. 4E).

Ancestral paralogs originating from the early vertebrate WGDs or older duplication events^16^ showed rod–cone expression partitioning (e.g., *gnat1* vs *gnat2*, *gnb1a/b* vs *gnb3b*, *pde6a/b* vs *pde6c*, *cnga1a/b* vs *cnga3a*, *saga* vs *arr3a/b*, *gucy2f* vs *gucy2d*, and *slc24a1* vs *slc24a2*). This expression partitioning is conserved across vertebrates and contributes to functional differences between rod and cone photoreceptors^49^. Divergent expression patterns were also observed for several teleost-specific paralogs, consistent with previous reports in zebrafish^49^. For example, *rgs9a* was expressed in cones whereas *rgs9b* was restricted to rods, while *gnb3a* was enriched in ON bipolar cells and *gnb3b* in cones (Fig. 3C,D). At the level of cone subtypes, *arr3a* was enriched in double cones (RH2 and LWS), whereas *arr3b* was restricted to single cones (SWS1), again consistent with observations in zebrafish^49,59^.

For most ohnolog pairs derived from the recent allopolyploidisation event, both copies were detected in the snRNA-seq dataset (Fig. 3D). Cases in which only one ohnolog was detected were generally attributable either to incomplete genome annotation (*gnb5a-A*, *pde6gb-B*, and *grk1b-A*) or to expression levels below the detection threshold (*rcvrnb-A*, *grk7b-A*, and *guca1ab.2-B*), as the corresponding duplicate was also expressed at very low levels. Opsins were an exception among phototransduction genes: consistent with the bulk RNA-seq data, ten opsin genes were detected in the adult snRNA-seq dataset, whereas four cone opsins (larval-specific *opn1sw1-B, opn1sw2-A, opn1mw3-A*, and transcriptionally inactive *opn1mw3-B*) were not detected. The detected ohnologs generally retained the cell-type expression patterns of their corresponding zebrafish orthologues, although a notable divergence was observed for *cngb1a*. In zebrafish, *cngb1a* is primarily expressed in rods^49^, whereas in the common barbel, the two *cngb1a* ohnologs showed complementary expression across photoreceptor types, with *cngb1a-A-1* enriched in cones and *cngb1a-B-1* predominantly expressed in rods (Fig. 3D), suggesting subfunctionalisation of this recent ohnolog pair. Several ohnolog pairs additionally displayed quantitative expression divergence, with one copy consistently expressed at higher levels than its duplicate across photoreceptor populations where both were detected (e.g., *gnb5b-B > gnb5b-A, saga-A-2 > saga-B-2, arr3a-B > arr3a-A, rcvrn2-A > rcvrn2-B*, and *rgs9a-B-1 > rgs9a-B-2*). Overall, expression divergence among retained ohnologs was predominantly quantitative, with differences in relative expression levels more common than complete partitioning of expression domains.

Additionally, to determine whether developmental switches similar to those observed for cone opsins extend to downstream components of the phototransduction cascade, we examined the ontogenetic expression patterns of two major gene families that directly interact with activated opsins: transducin alpha subunits (*gnat* genes) and arrestins, using bulk RNA-seq across developmental stages. Although the expression levels of these genes changed throughout ontogeny, we found no developmental switches comparable to those observed for cone opsins, nor evidence that the expression trajectories of individual phototransduction genes consistently tracked those of specific opsin paralogs (Supplementary Fig. S1E, Supplementary Table S3). However, *gnat1* (and, to a lesser extent, *gnat2*) showed a distinct developmental trend, with *gnat1-A* exhibiting higher relative expression during early developmental stages and the difference between the two ohnologs gradually diminishing over ontogeny (Supplementary Fig. S1E). Whether this pattern reflects functional differentiation between *gnat1* ohnologs or simply accompanies broader developmental changes in retinal gene regulation remains unclear.

Our results suggest that, following allopolyploidisation in the common barbel, opsins represent a particularly dynamic component of the phototransduction cascade, exhibiting pronounced expression divergence among retained duplicates. This pattern contrasts with observations from goldfish, another independently derived cyprinid allopolyploid, in which phototransduction and photoreceptor recovery genes, including opsins, generally retain highly similar expression patterns within ohnolog pairs^47^. Together, these findings show that the regulatory consequences of WGD are not uniform, even among independently derived allopolyploid lineages within the same family. More broadly, our findings highlight developmental regulation as a potential mechanism through which duplicated visual genes can acquire distinct expression roles and thereby be retained following polyploidisation, and emphasise the value of comparative analyses across independently derived polyploid lineages for understanding the evolutionary consequences of genome duplication.

## Methods

### Sampling

Retinae of larval and adult common barbel (*Barbus barbus*) were analysed in this study. Larval specimens were obtained through captive breeding at the Faculty of Fisheries and Protection of Waters in Vodňany, Czechia, following artificial reproduction for aquaculture purposes. Larvae were reared under controlled conditions at 20–21 °C, with daily water exchange to maintain stable water chemistry. Approximately 90% of the tank surface was covered with a dark cover throughout development to reduce light exposure. Larvae were sampled at 5, 6, 8, 14, 21, 28, and 42 days post-fertilisation (dpf), spanning major ontogenetic transitions from early benthic stages through a transient pelagic phase to a juvenile benthic lifestyle (i.e., hatching at 4 dpf, pelagic 11/12 - 28 dpf, and benthic from approximately 28 dpf).

Adult specimens were collected from wild populations by electrofishing at four river localities in the Czech Republic: the Berounka River (Liblín: 49°55′07.9″N, 13°32′43.9″E; Třímany: 49°56′16.7″N, 13°35′23.6″E), the Sázava River (Pikovice: 49°52′42.2″N, 14°25′37.1″E), and the Vltava River (Rájov: 48°51′04.2″N, 14°21′58.9″E). Sampling was conducted by authorised personnel under a permit issued by the Czech Anglers Union (Český rybářský svaz; CZ 03061). Fish were euthanised following established ethical guidelines. Whole larval specimens and dissected adult eyes were fixed and stored according to the requirements of subsequent analyses. A complete overview of all samples analysed in this study is provided in Supplementary Table S1.

### Genome annotation

We used two publicly available genome assemblies of the common barbel (*Barbus barbus*) from the NCBI assembly database. We first performed genome annotation using the fBarBab1.1 assembly (NCBI accession GCF_936440315.1), which was selected based on its higher quality and completeness. However, this assembly did not include information on chromosome pairs originating from the recent WGD event. We therefore used the ASM2356617v1 assembly (NCBI accession GCA_023566175.1) to transfer chromosome-pair information to fBarBab1.1. The ASM2356617v1 assembly identifies chromosome pairs using “A” and “B” labels (e.g., 1a and 1b); these labels distinguish chromosomes within a pair but do not consistently correspond to the two subgenomes. Chromosomes from ASM2356617v1 were aligned individually to fBarBab1.1, and chromosome names were transferred based on the best-scoring alignments. A table linking chromosome names across the two assemblies is provided in Supplementary Table S6.

Genome annotation was performed using an in-house annotation pipeline described by Kłodawska, Balart-García et al. (2026)^60^, which integrates short-read RNA-seq and protein homology evidence. RNA-seq evidence consisted of bulk retinal transcriptomes from multiple developmental stages of the common barbel generated in this study (Supplementary Table S1), together with a publicly available spleen transcriptome (NCBI accession SAMEA11296564). The predicted proteome of *Onychostoma macrolepi*s (NCBI accession GCF_012432095.1) served as protein homology evidence.

RNA-seq data were quality-filtered using fastp v0.22.0^61^ and aligned to the reference genome using HISAT2 v2.2.1^62^. To minimise potential mapping ambiguities associated with the tetraploid nature of the common barbel genome, each transcriptome sample was aligned independently to three versions of the reference genome: (1) a complete tetraploid reference genome, (2) a reference containing only A chromosomes and all unassigned short scaffolds, and (3) a reference containing only B chromosomes. All three reference genomes were soft-masked using a repeat library generated with RepeatModeler v1.0.11^63^ and subsequently applied with RepeatMasker v4.0.7 (www.repeatmasker.org). For each reference genome, SAM files generated from all RNA-seq samples were converted to BAM and concatenated using SAMtools v1.11^64^ to produce three final, reference-specific BAM datasets.

Downstream genome annotation employed StringTie v2.1.0^65^ and BRAKER3 v3.0.8^66^. StringTie was run independently using the BAM dataset generated against the complete reference genome and the BAM datasets generated against the separate A-chromosome and B-chromosome reference genomes. Multiple parameter combinations were evaluated to optimise transcript reconstruction. BRAKER3 was executed in two independent runs, both using the complete tetraploid reference genome. The first run used RNA-seq evidence derived from alignments to the complete reference genome, whereas the second run used RNA-seq evidence derived from the separate A-chromosome and B-chromosome reference genomes. Both runs incorporated protein homology evidence from the *Onychostoma macrolepis* proteome. The resulting GTF annotations were manually inspected in IGV v2.18.2^67^. For StringTie, the final annotation was generated from the complete-genome BAM dataset using stringent parameters (-c 2, -f 0.1, -m 300, -j 5, -a 30). For BRAKER3, the annotation generated using the separate A-chromosome and B-chromosome BAM datasets was selected for downstream analyses.

The selected StringTie and BRAKER3 annotations were subsequently merged and processed to generate a comprehensive, non-redundant genome annotation. This process followed the in-house pipeline^60^, with the *Onychostoma macrolepis* proteome used as protein homology evidence during coding sequence prediction. Mitochondrial genes were identified by manual inspection and added to the final genome annotation as custom GTF entries, with gene names prefixed by “MT-”. To distinguish genes originating from the recent WGD event, gene names were suffixed with “-A” or “-B” according to their position on A or B chromosomes respectively. Where duplicate gene names remained, additional suffixes were added to ensure that all gene names were unique.

### Phylogenetic analysis of the opsin genes

Phylogenetic analysis of opsin genes from the common barbel, common carp (*Cyprinus carpio*), and zebrafish (*Danio rerio*) was performed to investigate the evolutionary history of the opsin gene repertoire. Opsin gene sequences of the common barbel were extracted from the fBarBab1.1 genome assembly (NCBI accession GCF_936440315.1) according to the genome annotation generated in this study. Common carp opsin gene sequences, including pseudogenes, were retrieved from a published genome assembly (NCBI accession GCF_018340385.1). Zebrafish opsin gene sequences were downloaded from the NCBI GenBank database. Vertebrate ancient long opsin a (*valopa*) sequences from common carp and zebrafish, together with vertebrate ancient long opsin b (*valopb*) from zebrafish, were used as outgroups. All sequences included in the analysis and their corresponding accession numbers are listed in Supplementary Table S1.

Coding sequences of all opsin genes included in the dataset were processed in Geneious Prime® 2023.2.1 (www.geneious.com, Biomatters Ltd., Auckland, New Zealand). Multiple sequence alignment was performed using the MAFFT plugin (v1.5.0)^68^ implemented in Geneious Prime with default parameters. The resulting alignment was trimmed to exclude poorly alignable regions. Trimming was guided by the zebrafish LWS gene annotation, and the first and sixth exons were removed from the final alignment.

Phylogenetic analysis was performed using MrBayes v3.2.7a^69^. The GTR+I+Γ substitution model (nst = 6, rates = invgamma) was applied. Two independent Markov chain Monte Carlo (MCMC) runs were conducted, each with four chains and 10,000,000 generations. Trees and parameters were sampled every 1,000 generations. The initial 2,500 sampled trees from each run were discarded as burn-in, and the remaining samples were used to construct a majority-rule consensus tree with posterior probability support values. Convergence between runs was assessed by visual inspection of parameter traces in Tracer v1.7.2^70^. The final phylogenetic tree was visualised and edited in FigTree v1.4.4 (tree.bio.ed.ac.uk/software/figtree/).

### Opsin spectral sensitivity analysis

Absorption maxima ( λ_max_) of common barbel opsins were predicted using the Opsin Phenotype Tool for Inference of Color Sensitivity (OPTICS) v1.3^33,34^. Opsin protein sequences were generated by translating coding sequences identified in the fBarBab1.1 genome assembly (NCBI accession GCF_936440315.1) annotated in this study. Four OPTICS analyses were performed, evaluating all combinations of the whole-dataset and whole-dataset-MNM models with amino acid property and one-hot encoding methods, using bootstrap predictions in all cases (results shown in Supplementary Table S2). Predictions from the whole-dataset model with amino-acid property encoding were used as the primary estimates of λ_max_. OPTICS bootstrap visualisation plots were generated automatically from this analysis. Spectral sensitivity curves for individual opsins were reconstructed using a template proposed by Govardovskii et al. (2000)^35^.

### RNA isolation, cDNA library preparation, and sequencing

A total of 34 retinal transcriptomes were generated (Supplementary Table S1). Samples for RNA isolation were preserved in RNAlater and stored at −80 °C until processing. Larval eyes were used without further dissection, whereas adult retinae were carefully dissected immediately prior to RNA isolation. RNA was extracted from larval eyes using the QIAGEN RNeasy Micro Kit and from adult retinae using the QIAGEN RNeasy Mini Kit (QIAGEN, Hilden, Germany), following the respective manufacturer’s protocols. RNA concentration and quality were assessed using a Quantus fluorometer (Promega, Madison, WI, USA) and an Agilent 2100 Bioanalyzer (Agilent Technologies, Santa Clara, CA, USA), respectively. RNA isolates were stored at −80 °C until cDNA library preparation.

cDNA libraries were prepared using the NEBNext Ultra II Directional RNA Library Prep Kit for Illumina in combination with NEBNext Poly(A) mRNA Magnetic Isolation Module, NEBNext Multiplex Oligos for Illumina (New England Biolabs, Ipswich, MA, USA), and SPRIselect beads (Beckman Coulter, Brea, CA, USA). Library preparation was performed according to the manufacturer’s instructions using half-reaction volumes. Fragment size distribution in the libraries was assessed using an Agilent 2100 Bioanalyzer (Agilent Technologies, Santa Clara, CA, USA). If unwanted fragments were detected, additional size selection was performed using SPRIselect beads according to the manufacturer’s protocol.

Following library preparation and cleanup, library concentrations were quantified using a Quantus fluorometer (Promega, Madison, WI, USA), and libraries were pooled to achieve equimolar representation of each sample in the sequencing pool. Sequencing was outsourced to Novogene Co., Ltd. (for 8 dpf to adult stages) or Macrogen Europe B.V. (for 5 and 6 dpf stages).

### Bulk RNA-seq data analysis

Sequenced cDNA libraries were quality assessed using fastp^61^. Subsequent analyses were performed in Geneious Prime® 2023.2.1 (www.geneious.com, Biomatters Ltd., Auckland, New Zealand). For each sample, paired-end reads were mapped to reference sequences of opsin, arrestin, and transducin genes derived from the genome annotation generated in this study (Supplementary Data 1). Mapping was performed using the Geneious Mapper with custom sensitivity settings based on the Low Sensitivity/Fastest preset. The minimum read overlap was set to 50 bp, while all other mapping parameters were retained at their default values for the Low Sensitivity/Fastest preset. Only read pairs for which both reads mapped were retained, and reads with multiple equally good matches were assigned randomly.

Read counts mapped to each reference gene sequence were normalised for gene length and library size and expressed as reads per kilobase of transcript per million mapped reads (RPKM) (Supplementary Table S3). Relative expression of each opsin gene was calculated separately for cone and rod opsins and expressed as a percentage of total cone or rod opsin expression, respectively. For comparisons between cone types, the relative expression values of SWS1 and SWS2 opsins were summed to obtain total single-cone opsin expression, whereas the relative expression values of RH2 and LWS opsins were summed to obtain total double-cone opsin expression. Cone and rod opsin expression was compared as a percentage of total opsin expression. Results were visualised in R v4.5.0^71^ using the tidyverse package collection^72^, including ggplot2^73^. Visualisation was conducted within RStudio 2024.12.1+563^74^.

### Single-nucleus preparation, sorting, and sequencing

Two samples were prepared for single-nucleus RNA sequencing, yielding two independently generated and sequenced libraries (Supplementary Table S1). Dissected retinae of adult specimens were flash-frozen in liquid nitrogen and then stored at −80 °C before use. Nuclei were isolated according to the 10x Genomics Demonstrated Protocol (CG000366, Rev. D), with the following modifications: (i) In the final resuspension step of the nuclei isolation protocol, nuclei were resuspended in a resuspension buffer (1x PBS, 2% BSA, 0.2 U/μl RNase inhibitor) instead of Diluted Nuclei Buffer. (ii) A lower concentration of RNase inhibitor (0.1 U/μl ) was used in all other buffers (Wash Buffer, Lysis Buffer, Lysis Dilution Buffer). (iii) The total incubation time in the lysis step was 4 minutes. Isolated nuclei in the resuspension buffer were maintained on ice and transported for approximately 30 min to the Institute of Molecular Genetics of the Czech Academy of Sciences, where library preparation and sequencing were performed.

Nuclei from Sample 1 were stained with Hoechst 33342 and propidium iodide and sorted on a BD Influx Cell Sorter into chilled resuspension buffer. Propidium iodide was excited using a 561 nm laser and detected through a 610/20 nm bandpass filter. Hoechst was excited using a 355 nm UV laser and detected through a 455/50 nm bandpass filter. Following sorting, nuclei were concentrated by centrifugation at 600 × g for 5 minutes at 4 °C. Excess supernatant was discarded, and the nuclei pellet was resuspended in the remaining resuspension buffer. The final resuspension volume was adjusted based on the nuclei count obtained during sorting. Sample 2 was processed without fluorescence-activated nuclei sorting.

The quality and concentration of nuclei in both samples were manually assessed using a Leica DMi8 microscope following trypan blue staining. Single-nucleus RNA-seq libraries were prepared using the 10x Genomics Chromium Next GEM Single Cell 3′ Reagent Kit v3.1 (10x Genomics, Pleasanton, CA, USA) according to the manufacturer’s protocol (CG000315, Rev. A) on the 10x Genomics Chromium platform (Chromium Controller for Sample 1 and Chromium iX for Sample 2). The volume of nuclei suspension loaded into each reaction was calculated to achieve a target recovery of 6,000 nuclei per library. Library quality was assessed using an Agilent 2100 Bioanalyzer (Agilent Technologies, Santa Clara, CA, USA). Both libraries were sequenced on an Illumina NextSeq 2000 instrument (Illumina, San Diego, CA, USA). Sample 1 was sequenced using a NextSeq 500/550 High-Output v2.5 Kit (75 cycles) with a cDNA read length of 64 bp, and Sample 2 was sequenced using a NextSeq 1000/2000 P2 100 XLEAP-SBS Reagent Kit (100 Cycles) with a cDNA read length of 110 bp.

### Single-cell RNA-seq analysis

Both single-nucleus RNA-seq datasets generated from common barbel retinae were mapped to the fBarBab1.1 genome assembly (NCBI accession GCF_936440315.1) using the genome annotation generated in this study. Data processing and analysis were conducted following Kłodawska, Balart-García et al. (2026)^60^. Indexing and read quantification were carried out with SimpleAF^75^. The resulting count matrices were analysed using Seurat v5.3.0^76^ in R v4.5.0^71^, within RStudio 2024.12.1+563^74^.

Initial quality control involved filtering with thresholds determined separately for each sample based on preliminary inspection of quality metrics. For Sample 1, nuclei with fewer than 200 or more than 3,000 detected genes (nFeature_RNA), mitochondrial transcript content exceeding 1%, more than 15,000 transcript counts (nCount_RNA), or low transcript complexity (log_10_ genes per UMI ≤ 0.93) were excluded from further analysis. For Sample 2, nuclei with fewer than 200 or more than 2,500 detected genes, mitochondrial transcript content exceeding 2%, more than 10,000 transcript counts, or low transcript complexity (log_10_ genes per UMI ≤ 0.93) were excluded. Ambient RNA contamination and putative doublets were detected and removed using scCDC^77^ and DoubletFinder^78^, respectively. Following all these processing steps, Sample 1 contained 4,670 nuclei with expression detected for 35,256 genes, while Sample 2 contained 5,888 nuclei with expression detected for 34,015 genes, indicating comparable levels of gene detection.

Corrected singlet datasets were integrated using Seurat’s anchor-based integration workflow with SCTransform normalisation. Following dimensionality reduction using principal component analysis (PCA), clustering was performed using Seurat’s FindNeighbors and FindClusters functions. The resulting clusters were visualised in a UMAP embedding and displayed using DimPlot, and the representation of both datasets across cell populations was assessed. Subsequently, differential gene expression (DGE) analysis was performed using Seurat’s FindMarkers function. Cluster identities were assigned based on differentially expressed genes and validated using known retinal marker genes from the literature^13,47,49^ and the DanioCell database^46^. Clusters corresponding to non-retinal cell types were excluded. A population of microglial cells was also removed because it was dispersed across multiple retinal cell clusters in the UMAP rather than forming a discrete cluster. The PCA, clustering, DGE, and annotation steps were then repeated on the filtered dataset to generate the final integrated and annotated atlas of retinal cells in the common barbel.

Cell type annotation was based on differentially expressed genes and informed by published marker genes^13,47,49^ as well as the DanioCell database^46^ (Supplementary Table S4). While several clusters could be assigned unambiguously based on highly specific marker genes (e.g., *rho* in rods and *mag* in oligodendrocytes), others exhibited overlapping expression profiles that precluded assignment based on a single diagnostic marker. In particular, retinal interneuron and ganglion cell clusters proved difficult to annotate, partly because of their low representation in the dataset. Consequently, these annotations were based on the combined expression of multiple marker genes. One amacrine cell population (ACs-1) lacked specific diagnostic markers but was assigned based on its overall transcriptional profile, which was consistent with an inhibitory neuronal cell type characterised by abundant synaptic machinery and diverse ion channels. Another cluster, designated “Neurons”, displayed a neuronal transcriptional signature but could not be confidently assigned to a specific neuronal cell type. Expression of representative marker genes enriched in individual cell type populations was visualised with Seurat’s DotPlot. The relative proportions of major retinal cell types recovered from each of the two common barbel samples were visualised as donut charts using the ggplot2 package^73^.

To further investigate gene expression among cone subtypes, the cone cluster was subsetted and reanalysed using PCA, clustering, and UMAP. Clusters were visualised using Seurat’s DimPlot, and marker gene expression was examined using FeaturePlot. Three subclusters corresponding to the SWS1 (UV, PR4), RH2 (green, PR2), and LWS (red, PR1) cone subtypes were identified based on subtype-specific marker genes^49–51^, with opsin expression providing additional support for their assignment. DGE analysis was then performed between cone subclusters, and the top 10 marker genes for each cone subtype were visualised using DotPlot.

Additional DotPlots were generated to examine expression of phototransduction and photoreceptor recovery genes. A list of 103 phototransduction and photoreceptor recovery genes was compiled based on published literature^16,49^. We analysed their expression across photoreceptor clusters (rods, rod precursors, and cones) in the integrated dataset. Genes expressed in cone photoreceptors were further examined across the three cone subtypes within the subsetted cone dataset.

### Fluorescence in situ hybridisation

Fluorescence in situ hybridisation (FISH) was performed on whole-mounted adult retinae to localise cone opsin expression, following the protocol described in Musilova et al. (2019)^79^, with minor modifications.

RNA extracted from the adult retina was used for cDNA synthesis using the Mint-2 cDNA Synthesis Kit (Evrogen, Moscow, Russia). Probe templates were amplified with primers designed to target regions of sequence divergence between opsin ohnologs. Amplified fragments were approximately 450–500 bp in length and spanned exons 1 to 3 for RH2 genes, exons 1 to 4 for SWS1 genes, exon 3 to the 3′ untranslated region (UTR) for SWS2 genes, and exon 5 to the 3′ UTR for LWS genes. PCR products were purified from agarose gels using the Gel/PCR DNA Fragments Extraction Kit (Geneaid Biotech Ltd., New Taipei City, Taiwan). Fluorescein (FL)- and digoxigenin (DIG)-labelled RNA probes were synthesised from the purified DNA templates using T3 RNA polymerase.

Eye samples used for FISH were fixed with 4% paraformaldehyde (PFA) for 30 min, transferred to 100% methanol, and stored at –20 °C. Prior to hybridisation, retinae were dissected and their orientation was marked with a dorsal incision. Each retina was hybridised with a pair of selected opsin probes. The following probe combinations were tested: *sw1-A* and *sw2-B*, *mw4-A* and *lw1-B*, *lw1-A* and *lw1-B*, and *mw4-A* and *mw4-B*. Probe signals were amplified using Tyramide SuperBoost™ Kits with Alexa Fluor™ 488 Tyramide for FL-labelled probes and Alexa Fluor™ 594 Tyramide for DIG-labelled probes (Invitrogen, Thermo Fisher Scientific, Waltham, MA, USA). Finally, retinae were transferred to 70% glycerol and mounted as whole mounts on microscope slides with the photoreceptor layer facing downward.

Imaging was performed on a Zeiss LSM 880 laser scanning confocal microscope at the Viničná Microscopy Core Facility, and images were processed using Fiji^80^.

## Acknowledgements

We would like to thank Dr. Miroslav Švátora, and Ing. Hynek Dort from the Czech Anglers Union for their assistance during collections in the field. We are grateful to Anna Nováková for her laboratory support with fluorescence in situ labelling and single nucleus isolation. We also thank the Laboratory of Flow Cytometry at the Institute of Molecular Genetics of the Czech Academy of Sciences for nuclei sorting, and Dr. Michal Kolář and his team at the same institute for their assistance and expertise in single-nucleus RNA sequencing.

The study was supported by the Grant Agency of Charles University (GAUK for ZK; project no. 350122), the PRIMUS research programme of the Charles University (for ZM), the Czech Science Foundation (GACR for ZM; project no. 21-31712S, “Zoom in the fish eye & blood: molecular evolution of functional adaptations in deep-sea and freshwater fishes”), and the European Research Council (ERC) Consolidator grant (for ZM; project no. 101122542, “SensingDEEP”).

Measurements using a Quantus fluorometer and an Agilent 2100 Bioanalyzer were performed at the Laboratory OMICS - Genomics in BIOCEV. Microscopy was performed at the Vinicna Microscopy Core Facility (RRID:SCR_026602) co-financed by the Czech-BioImaging large RI project LM2023050. Computational resources were provided by the e-INFRA CZ project (ID:90254), supported by the Ministry of Education, Youth and Sports of the Czech Republic.

## Data availability

Single-nucleus RNA-seq and bulk RNA-seq data have been deposited in the NCBI BioProject PRJNA1521124. All custom scripts used in this study are available in GitHub: https://github.com/FishEvoLab/Barbus_project.

## Author contributions

Z.K. and Z.M. conceived the study. Z.K., V.T., and Z.M. collected samples in the field. P.L. provided samples of larval stages. V.T. and Z.K. processed samples for bulk RNA-seq. Z.K. performed bulk transcriptome analysis. Z.K. processed samples for single-nucleus RNA-seq. D.B. performed genome masking and assisted during genome annotation. P.B.G. developed single-cell RNA-seq analysis pipeline. Z.K. performed the genome annotation and single-cell RNA-seq analyses. Z.K. performed the FISH experiments. Z.K. performed the phylogenetic analysis. Z.M. predicted opsin spectral sensitivities with OPTICS. Z.K. and Z.M. drafted the manuscript, figures, tables, and supplementary material. Z.M. supervised this study. All the coauthors contributed to the final version of the manuscript.

## Supplementary information for

**Supplementary Figure S1:**
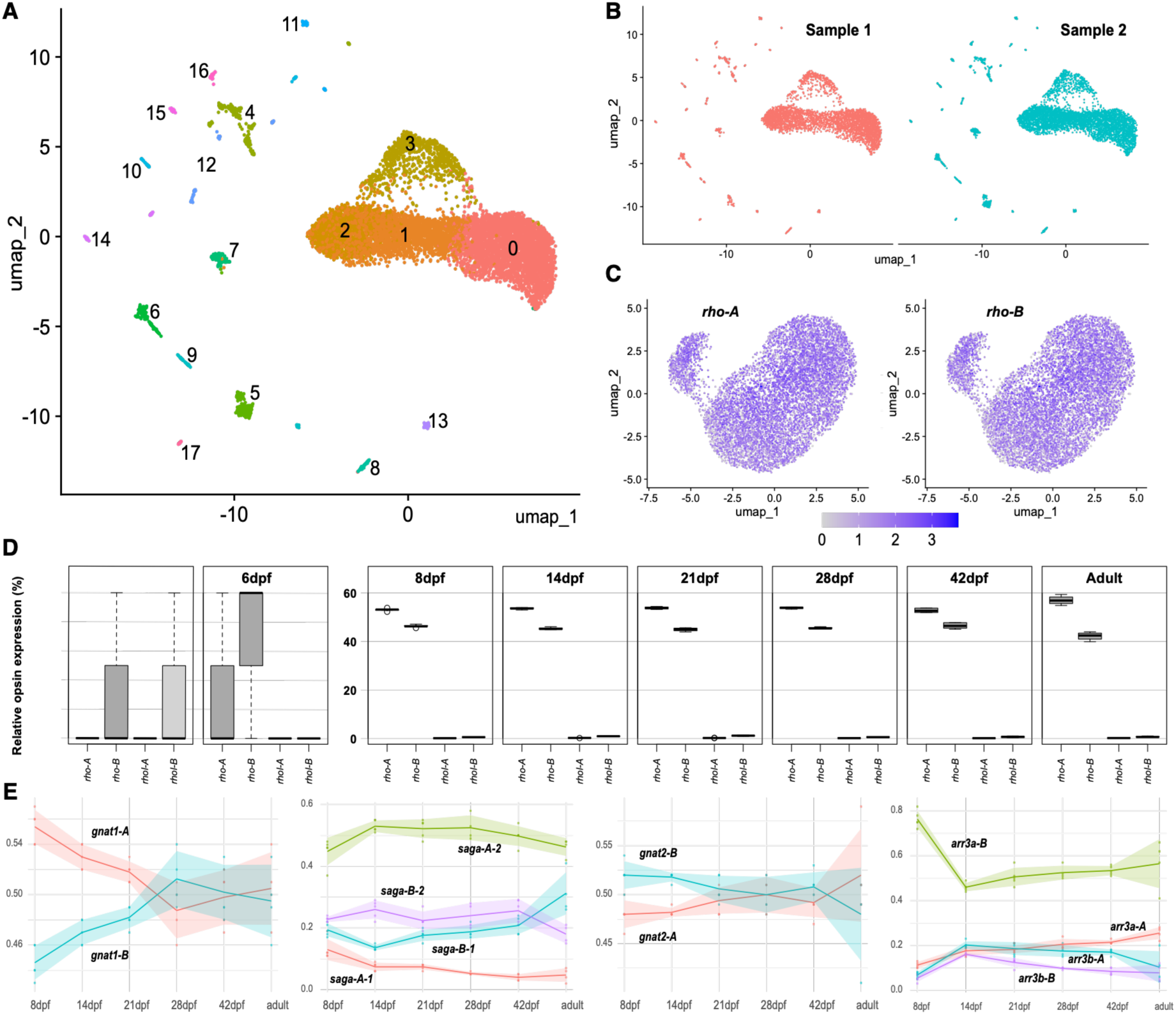
Single-nucleus clustering and developmental changes in expression of selected genes in the common barbel retina. (A) UMAP visualisation of the integrated snRNA-seq dataset showing all clusters before retinal cell type annotation. (B) UMAP visualisation of nuclei from the two common barbel samples profiled by single-nucleus sequencing: Sample 1 (nuclei sorted prior library construction) and sample 2 (unsorted nuclei). All clusters were represented in both samples. (C) Expression of rhodopsin ohnologs (*rho-A* and *rho-B*) in a separately visualised rod cluster. Rhodopsin ohnologs were co-expressed across the rod population. (D) Developmental series of rod opsin expression in the common barbel based on bulk retinal transcriptomes. Relative expression of rhodopsin (*rho*) and rhodopsin-like (*rhol*) genes is shown as box plots, with each opsin represented as a proportion of total rod opsin expression. The “-A” and “-B” suffixes denote respective ohnologs. Note the different y-axis scales. (E) Developmental changes in transducin and arrestin expression based on bulk retinal transcriptomes. Lines represent mean expression levels, and ribbons show standard deviations. The “-A” and “-B” suffixes denote respective ohnologs.

### Captions for Supplementary Tables

**Supplementary Table S1.** List of samples and datasets used in this study, including bulk RNA-seq data and snRNA-seq data.

**Supplementary Table S2.** Opsin spectral sensitivity predictions from OPTICS^33,34^. The table shows predictions for all combinations of the whole-dataset and whole-dataset-MNM models with amino acid property and one-hot encoding methods.

**Supplementary Table S3.** Results of bulk RNA-seq data mapping to reference gene sequences of opsins, arrestins, and transducins. The table includes raw read counts for all samples and read counts normalised for gene length and library size (reads per kilobase of transcript per million mapped reads, RPKM).

**Supplementary Table S4.** Differentially expressed genes used for retinal cell cluster annotation in the snRNA-seq dataset. The table reports DGE results for each cluster, including gene name, P value, average log₂ fold change, adjusted P value, percentage of expressing cells (pct.1 and pct.2), cluster number, and assigned cell type.

**Supplementary Table S5.** Differential gene expression analysis of cone subclusters within the cone cluster in the common barbel retina. The table lists the top 10 differentially expressed genes for each cone subcluster, including gene name, P value, average log₂ fold change, adjusted P value, percentage of expressing cells (pct.1 and pct.2), and assigned cone subtype (cluster_plot: LWS, SWS1, or RH2).

**Supplementary Table S6.** Conversion of chromosome names and accession numbers between the two published common barbel genome assemblies and the chromosome nomenclature used in this study.

### Caption for Supplementary Data 1

**Supplementary Data 1**. Coding sequences of opsin, arrestin, and transducin genes derived from the genome annotation generated in this study and used as references for mapping bulk RNA-seq data. Sequences are provided in FASTA format.

## References

1. Wertheim, B., Beukeboom, L. W. & van de Zande, L. Polyploidy in animals: Effects of gene expression on sex determination, evolution and ecology. Cytogenet. Genome Res. 140, 256–269 (2013).

2. De Smet, R. & Van de Peer, Y. Redundancy and rewiring of genetic networks following genome-wide duplication events. Curr. Opin. Plant Biol. 15, 168–176 (2012).

3. Van de Peer, Y., Maere, S. & Meyer, A. The evolutionary significance of ancient genome duplications. Nat. Rev. Genet. 10, 725–732 (2009).

4. Meyer, A. & Van de Peer, Y. From 2R to 3R: Evidence for a fish-specific genome duplication (FSGD). BioEssays 27, 937–945 (2005).

5. Dehal, P. & Boore, J. L. Two Rounds of Whole Genome Duplication in the Ancestral Vertebrate. PLoS Biol. 3, 1700–1708 (2005).

6. Luo, J. et al. Tempo and mode of recurrent polyploidization in the *Carassius auratus* species complex (Cypriniformes, Cyprinidae). Heredity (Edinb). 112, 415–427 (2014).

7. Pasquier, J. et al. Evolution of gene expression after whole-genome duplication: New insights from the spotted gar genome. J. Exp. Zool. B Mol. Dev. Evol. 328, 709–721 (2017).

8. Van de Peer, Y., Mizrachi, E. & Marchal, K. The evolutionary significance of polyploidy. Nat. Rev. Genet. 18, 411–424 (2017).

9. Mable, B. K., Alexandrou, M. A. & Taylor, M. I. Genome duplication in amphibians and fish: An extended synthesis. J. Zool. 284, 151–182 (2011).

10. Soltis, P. S., Marchant, D. B., Van de Peer, Y. & Soltis, D. E. Polyploidy and genome evolution in plants. Curr. Opin. Genet. Dev. 35, 119–125 (2015).

11. Xie, C. et al. Chromosomal fusions trigger rediploidization of autopolyploid genomes. Nature 654, 706–713 (2026).

12. Yang, L., Naylor, G. J. P. & Mayden, R. L. Deciphering reticulate evolution of the largest group of polyploid vertebrates, the subfamily cyprininae (Teleostei: Cypriniformes). Mol. Phylogenet. Evol. 166, 107323 (2022).

13. Hahn, J. et al. Evolution of neuronal cell classes and types in the vertebrate retina. Nature 624, 415–424 (2023).

14. Baker, S. A. & Kerov, V. Photoreceptor inner and outer segments. in Current Topics in Membranes vol. 72, 231–265 (Elsevier, 2013).

15. Masland, R. H. The Neuronal Organization of the Retina. Neuron 76, 266–280 (2012).

16. Lamb, T. D. Evolution of the genes mediating phototransduction in rod and cone photoreceptors. Prog. Retin. Eye Res. 76, 100823 (2020).

17. Yokoyama, S. Molecular evolution of vertebrate visual pigments. Prog. Retin. Eye Res. 19, 385– 419 (2000).

18. Yokoyama, S. Evolution of Dim-Light and Color Vision Pigments. Annu. Rev. Genomics Hum. Genet. 9, 259–282 (2008).

19. Carleton, K. L., Escobar-Camacho, D., Stieb, S. M., Cortesi, F. & Marshall, N. J. Seeing the rainbow: mechanisms underlying spectral sensitivity in teleost fishes. J. Exp. Biol. 223, jeb193334 (2020).

20. Musilova, Z., Salzburger, W. & Cortesi, F. The Visual Opsin Gene Repertoires of Teleost Fishes: Evolution, Ecology, and Function. Annu. Rev. Cell Dev. Biol. 37, 441–468 (2021).

21. Evans, B. I. & Fernald, R. D. Metamorphosis and fish vision. J. Neurobiol. 21, 1037–1052 (1990).

22. Allison, W. T. et al. Ontogeny of Cone Photoreceptor Mosaics in Zebrafish. J. Comp. Neurol. 518, 4182–4195 (2010).

23. Nandamuri, S. P., Yourick, M. R. & Carleton, K. L. Adult plasticity in African cichlids: Rapid changes in opsin expression in response to environmental light differences. Mol. Ecol. 26, 6036– 6052 (2017).

24. Chang, C. H. et al. Sequence Analysis and Ontogenetic Expression Patterns of Cone Opsin Genes in the Bluefin Killifish (*Lucania goodei*). J. Hered. 112, 357–366 (2021).

25. Shand, J. et al. The influence of ontogeny and light environment on the expression of visual pigment opsins in the retina of the black bream, *Acanthopagrus butcheri*. J. Exp. Biol. 211, 1495–1503 (2008).

26. Cortesi, F. et al. Ancestral duplications and highly dynamic opsin gene evolution in percomorph fishes. Proc. Natl. Acad. Sci. U. S. A. 112, 1493–1498 (2015).

27. Wood, P. & Partridge, J. C. Opsin substitution induced in retinal rods of the eel (*Anguilla anguilla* (L.)): A model for G-protein-linked receptors. Proc. R. Soc. B 254, 227–232 (1993).

28. Spady, T. C. et al. Evolution of the cichlid visual palette through ontogenetic subfunctionalization of the opsin gene arrays. Mol. Biol. Evol. 23, 1538–1547 (2006).

29. Chen, J. N., Samadi, S. & Chen, W. J. Rhodopsin gene evolution in early teleost fishes. PLoS One 13, e0206918 (2018).

30. Musilova, Z. et al. Vision using multiple distinct rod opsins in deep-sea fishes. Science 364, 588–592 (2019).

31. Policarpo, M., Fogg, L. G., Cortesi, F. & Salzburger, W. Evolution of the Nonvisual and Visual Opsin Gene Repertoire in Ray-Finned Fishes. Genome Biol. Evol. 17, evaf129 (2025).

32. Cortesi, F. et al. From crypsis to mimicry: Changes in colour and the configuration of the visual system during ontogenetic habitat transitions in a coral reef fish. J. Exp. Biol. 219, 2545–2558 (2016).

33. Frazer, S. A. & Oakley, T. H. Accessible and robust machine learning approaches to improve the opsin genotype-phenotype map. Mol. Biol. Evol. 43, msag138 (2026).

34. Frazer, S. A., Baghbanzadeh, M., Rahnavard, A., Crandall, K. A. & Oakley, T. H. Discovering genotype–phenotype relationships with machine learning and the Visual Physiology Opsin Database (VPOD). GigaScience 13, giae073 (2024).

35. Govardovskii, V. I., Fyhrquist, N., Reuter, T., Kuzmin, D. G. & Donner, K. In search of the visual pigment template. Vis. Neurosci. 17, 509–528 (2000).

36. Cheng, C. L. & Flamarique, I. N. Chromatic organization of cone photoreceptors in the retina of rainbow trout: single cones irreversibly switch from UV (SWS1) to blue (SWS2) light sensitive opsin during natural development. J. Exp. Biol. 210, 4123–4135 (2007).

37. Lupše, N. et al. Developmental changes of opsin gene expression in ray-finned fishes (Actinopterygii). Proc. R. Soc. B 289, 20221855 (2022).

38. Britt, L. L., Loew, E. R. & McFarland, W. N. Visual pigments in the early life stages of Pacific northwest marine fishes. J. Exp. Biol. 204, 2581–2587 (2001).

39. Johnsen, S. & Widder, E. A. Ultraviolet absorption in transparent zooplankton and its implications for depth distribution and visual predation. Mar. Biol. 138, 717–730 (2001).

40. Nowosad, J., Kupren, K., Biegaj, M. & Kucharczyk, D. Allometric and ontogenetic larval development of common barbel during rearing under optimal conditions. Animal 15, 100107 (2021).

41. Vilizzi, L. & Copp, G. H. Interstitial movement and emergence of barbel *Barbus barbus* free embryos and larvae. J. Fish Biol. 82, 1057–1063 (2013).

42. Bischoff, A. & Freyhof, J. Seasonal shifts in day-time resource use of 0+ barbel, *Barbus barbus*. Environ. Biol. Fishes 56, 199–212 (1999).

43. Lupše, N. et al. Visual Gene Expression Reveals a cone-to-rod Developmental Progression in Deep-Sea Fishes. Mol. Biol. Evol. 38, 5664–5677 (2021).

44. Schweikert, L. E. & Grace, M. S. Spectral Sensitivity Change May Precede Habitat Shift in the Developing Retina of the Atlantic Tarpon (*Megalops atlanticus*). Physiol. Biochem. Zool. 90, 553–563 (2017).

45. Bowmaker, J. K., Semo, M., Hunt, D. M. & Jeffery, G. Eel visual pigments revisited: The fate of retinal cones during metamorphosis. Vis. Neurosci. 25, 249–255 (2008).

46. Sur, A. et al. Single-cell analysis of shared signatures and transcriptional diversity during zebrafish development. Dev. Cell 58, 3028–3047.e12 (2023).

47. Kon, T. et al. Single-cell transcriptomics of the goldfish retina reveals genetic divergence in the asymmetrically evolved subgenomes after allotetraploidization. *Commun*. Biol. 5, 1404 (2022).

48. Li, Z. et al. Cross-species single-cell transcriptomic atlas of retinal photoreceptors reveals molecular signatures underlying color vision adaptation. Cell Rep. 44, 116164 (2025).

49. Ogawa, Y. & Corbo, J. C. Partitioning of gene expression among zebrafish photoreceptor subtypes. Sci. Rep. 11, 17340 (2021).

50. Baden, T. et al. A standardized nomenclature for the rods and cones of the vertebrate retina. PLoS Biol. 23, e3003157 (2025).

51. Tommasini, D., Yoshimatsu, T., Puthussery, T., Baden, T. & Shekhar, K. Comparative transcriptomic insights into the evolution of vertebrate photoreceptor types. Curr. Biol. 35, 2228–2239.e4 (2025).

52. Compère, P. & Poncin, P. Microscopic observation of the retinal photoreceptor layer of the common barbel (Teleostei: Cyprinidae). Belg. J. Zool. 126, 49–56 (1996).

53. Stell, W. K. & Hárosi, F. I. Cone structure and visual pigment content in the retina of the goldfish. Vision Res. 16, 647–657 (1976).

54. Bowmaker, J. K. & Kunz, Y. W. Ultraviolet receptors, tetrachromatic colour vision and retinal mosaics in the brown trout (*Salmo trutta*): Age-dependent changes. Vision Res. 27, 2101–2108 (1987).

55. Marc, R. E. & Sperling, H. G. The chromatic organization of the goldfish cone mosaic. Vision Res. 16, 1211–1224 (1976).

56. Tohya, S., Mochizuki, A. & Iwasa, Y. Difference in the Retinal Cone Mosaic Pattern Between Zebrafish and Medaka: Cell-Rearrangement Model. J. Theor. Biol. 221, 289–300 (2003).

57. Takechi, M. & Kawamura, S. Temporal and spatial changes in the expression pattern of multiple red and green subtype opsin genes during zebrafish development. J. Exp. Biol. 208, 1337–1345 (2005).

58. Dalton, B. E., Loew, E. R., Cronin, T. W. & Carleton, K. L. Spectral tuning by opsin coexpression in retinal regions that view different parts of the visual field. Proc. R. Soc. B 281, 20141980 (2014).

59. Renninger, S. L., Gesemann, M. & Neuhauss, S. C. F. Cone arrestin confers cone vision of high temporal resolution in zebrafish larvae. Eur. J. Neurosci. 33, 658–667 (2011).

60. Kłodawska, M. et al. Rapid evolution of cone identity and retinal topography in deep-water crater lake cichlids. Preprint at bioRxiv 10.64898/2026.09.16.751957 (2026).

61. Chen, S., Zhou, Y., Chen, Y. & Gu, J. fastp: an ultra-fast all-in-one FASTQ preprocessor. Bioinformatics 34, i884–i890 (2018).

62. Kim, D., Paggi, J. M., Park, C., Bennett, C. & Salzberg, S. L. Graph-based genome alignment and genotyping with HISAT2 and HISAT-genotype. Nat. Biotechnol. 37, 907–915 (2019).

63. Flynn, J. M. et al. RepeatModeler2 for automated genomic discovery of transposable element families. Proc. Natl. Acad. Sci. U. S. A. 117, 9451–9457 (2020).

64. Danecek, P. et al. Twelve years of SAMtools and BCFtools. Gigascience 10, giab008 (2021).

65. Shumate, A., Wong, B., Pertea, G. & Pertea, M. Improved transcriptome assembly using a hybrid of long and short reads with StringTie. PLoS Comput. Biol. 18, e1009730 (2022).

66. Gabriel, L. et al. BRAKER3: Fully automated genome annotation using RNA-seq and protein evidence with GeneMark-ETP, AUGUSTUS, and TSEBRA. Genome Res. 34, 769–777 (2024).

67. Robinson, J. T. et al. Integrative genomics viewer. Nat. Biotechnol. 29, 24–26 (2011).

68. Katoh, K. & Standley, D. M. MAFFT Multiple Sequence Alignment Software Version 7: Improvements in Performance and Usability. Mol. Biol. Evol. 30, 772–780 (2013).

69. Ronquist, F. et al. MrBayes 3.2: Efficient Bayesian Phylogenetic Inference and Model Choice Across a Large Model Space. Syst. Biol. 61, 539–542 (2012).

70. Rambaut, A., Drummond, A. J., Xie, D., Baele, G. & Suchard, M. A. Posterior Summarization in Bayesian Phylogenetics Using Tracer 1.7. Syst. Biol. 67, 901–904 (2018).

71. R Core Team. R: A Language and Environment for Statistical Computing (R Foundation for Statistical Computing, 2025); https://www.r-project.org/

72. Wickham, H. et al. Welcome to the Tidyverse. J. Open Source Softw. 4, 1686 (2019).

73. Wickham, H. ggplot2: Elegant Graphics for Data Analysis (Springer, 2016).

74. Posit team. RStudio: Integrated Development Environment for R (Posit Software, PBC, 2025); https://posit.co/

75. He, D. & Patro, R. simpleaf: a simple, flexible, and scalable framework for single-cell data processing using alevin-fry. Bioinformatics 39, btad614 (2023).

76. Hao, Y. et al. Dictionary learning for integrative, multimodal and scalable single-cell analysis. Nat. Biotechnol. 42, 293–304 (2024).

77. Wang, W. et al. scCDC: a computational method for gene-specific contamination detection and correction in single-cell and single-nucleus RNA-seq data. Genome Biol. 25, 136 (2024).

78. McGinnis, C. S., Murrow, L. M. & Gartner, Z. J. DoubletFinder: Doublet Detection in Single-Cell RNA Sequencing Data Using Artificial Nearest Neighbors. Cell Syst. 8, 329–337.e4 (2019).

79. Musilova, Z. et al. Evolution of the visual sensory system in cichlid fishes from crater lake Barombi Mbo in Cameroon. Mol. Ecol. 28, 5010–5031 (2019).

80. Schindelin, J. et al. Fiji: an open-source platform for biological-image analysis. Nat. Methods 9, 676–682 (2012).

